# A genome-wide association study identified new variants associated with mathematical abilities in Chinese children

**DOI:** 10.1101/2022.10.23.513388

**Authors:** Liming Zhang, Zhengjun Wang, Zijian Zhu, Qing Yang, Chen Cheng, Shunan Zhao, Chunyu Liu, Jingjing Zhao

**Affiliations:** School of Psychology, Shaanxi Normal University, and Shaanxi Provincial Key Research Center of Child Mental and Behavioral Health, Xi’an, China; Department of Psychiatry, SUNY Upstate Medical University, Syracuse, NY, USA

**Keywords:** Genome-wide association study, quantitative trait, mathematical ability, enrichment

## Abstract

Mathematical ability is moderately heritable, and it is a complex trait which can be evaluated in several different categories. A few genetic studies have been published on general mathematical ability. However, no genetic study focused on specific mathematical ability categories. In this study, we separately performed genome-wide association studies (GWASs) on 11 mathematical ability categories in 1 146 students from Chinese elementary schools. We identified 7 genome-wide significant single nucleotide polymorphisms (SNPs) with strong linkage disequilibrium (LD) among each other (all r^2^>0.8) associated with mathematical reasoning ability (top SNP: rs34034296, *p*=2.01×10^−8^, nearest gene: CUB and Sushi multiple domains 3, CSMD3). We replicated one SNP (rs133885) from 585 SNPs previously reported to be associated with general mathematical ability associated with division ability in our data (*p*=1.053×10^−5^). In the gene- and gene-set enrichment analysis by MAGMA, we found three significant enrichments of associations with three mathematical ability categories for three genes. We also observed four significant enrichments of associations with four mathematical ability categories for three gene sets. Our results suggest new candidate genetic loci for the genetics of mathematical ability.

## INTRODUCTION

Mathematical ability is a complex trait which is the key to excellent performance in study and work. Substantial heritability estimates (0.2-0.9) have been reported in previous studies indicating that mathematical ability was influenced by genetic factors[1–5]. In recent years, several GWASs have been applied to study genetic components associated with mathematical ability [6–11]. Among these GWASs, rs133885 was considered to be associated with mathematical ability (*n*=699, *p*=7.71×10^−10^) [11], but the finding was not replicated in another study [12]. Then, researchers found four loci associated with Chinese students’ curriculum scores of mathematics by conducting a meta-analysis [7]. Later, Lee and colleagues performed a GWAS on two phenotypes reflecting mathematical ability in large samples (*n_1_*=564 698, *n_2_*=430 445), they found a total of 983 loci at genome-wide significant level [10].

According to the Syllabus of Primary and Secondary Schools in China, mathematical ability can be divided into specific categories such as calculation ability (addition, subtraction, multiplication, division, etc.), logical reasoning ability, spatial ability and applied mathematics ability. According to UK National Curriculum (NC) Key Stage 2 criteria, the mathematical knowledge evaluated by the examination include number (number and position value, addition, subtraction, multiplication, division, etc.), measurement, geometry (properties of shapes, position and direction) and statistics. These mathematical ability categories are usually measured by some specific scales. The Heidelberg mathematics test is a well-established test with good reliability and validity, which considered to be one of the most comprehensive tests to measure different mathematical ability categories [13].

Previous GWASs mostly used composite scores as phenotypes (self-reported math ability, NC-based teacher ratings, NC-based web-based testing, academic achievement, etc.), except for one study, which used test measurement (numerosity judgements), a composite score of two tests (addition and multiplication) and a composite score of three tests (addition, multiplication and numerosity judgements) as three phenotypes, respectively [11]. To our knowledge, no researchers have systematically used a wide range of mathematical tests, the detailed subphenotypes such as those in Heidelberg mathematics test, to screen for genetic associations. In addition, none of past large scale genetic association studies of mathematical abilities have used asian populations.

In the present study, we performed a GWAS in students of the Chinese elementary school to discover the genetic basis of mathematical ability measured by Heidelberg mathematics test [13]. We separately analyzed eleven mathematical ability categories. Then, we also tested 585 SNPs that have been reported to be associated with general mathematical ability in our data for the eleven categories. Finally, we found some significant enrichment of associations with categories for specific genes or gene sets.

## METHODS AND MATERIALS

### Participants

A cohort of primary school students from two provinces in the northwestern part of China, including Shaanxi province and Gansu province, were recruited as participants in this study. Nonverbal intelligences of these children were measured by Raven’s Standard Progressive Matrices individually [14]. According to Zhang’s local norms [15], all children had normal IQ. In total, 1 182 participants were eligible for subsequent genotyping and association analysis. This study was approved by the ethics committee of Shaanxi Normal University, and written informed consent was obtained for all the participants’ parents.

### Phenotypic measure

A Chinese version of Heidelberg mathematics test was used to test the mathematical ability for the Chinese cohort [13–16]. The test included 11 subtests corresponding to 11 mathematical ability categories divided into two areas, namely the arithmetic operations area and the numerical-logical and spatial-visual skills area. Area of arithmetic operations assess performance in six tasks, including addition, subtraction, multiplication, division, equation and magnitude perception tasks. Area of numeric-logical and spatial-visual evaluate performances in five tasks, including mathematical reasoning, visual size estimation, spatial conception, quantity counting and visuomotor tasks. A brief description of this test is shown in Table S1.

The test-retest reliabilities of the whole test and the area tests were r = .69-.89 and r = .87-.93, respectively [13]. Later investigations revealed that correlations between whole test score and mathematical curriculum scores were r = .67 and r = .68 respectively in two different samples [17]. The split-half reliability of the Chinese version of Heidelberg mathematics test was 0.9 and the test score was also correlated with Chinese students’ curriculum scores of mathematics [16].

Students were tested in groups. Each subtest was required to complete in a given amount of time. Testing process of this test is shown in the supplementary methods. All analyses were based on the raw data.

Scores of subtests showed moderate to high degree of correlations with each other (see Table S2).

### Genotype quality control and imputation

Genomic DNAs were extracted from saliva samples by Hi-Swab DNA Kit (Lot#R7220, Cat.#DP362-02) of Tiangen Biotechnology in Beijing, China. Genome-wide genotyping experiments used Illumina Asian screening array (650K) supplied by Compass Biotechnology in Beijing, China. There were 1 177 samples and 704 415 SNPs in the original data. Genotype quality control was conducted in PLINK v1.90. SNPs were removed if they displayed a Hardy-Weinberg Equilibrium (HWE) <10^−5^, a minor allele frequency (MAF) <0.01, or a variant call rate <0.95. Individuals were removed if they displayed a sex discrepancy between the records and the genetically inferred data [18, 19], an unexpected duplicates or probable relatives (PI-HAT>0.20), or a genotyping call rate <0.9. After this step of quality control, 1 146 samples and 497 823 SNPs were left.

Imputation was conducted using the Genome Asia Pilot-GasP (GRCh37/hg19) reference panel on the Michigan Imputation Server (Minimac4). After imputation, there were 8 625 058 SNPs. Imputed SNPs were removed if they showed a R^2^ <0.6. After the imputation, we carried out a quality control again. SNPs were removed if they displayed a HWE <10^−5^, a MAF <0.05, or a variant call rate <0.95. Individuals were removed if they displayed a genotyping call rate <0.9. Finally, there were 5 406 859 SNPs and 1 146 samples left.

### Statistical analysis

After imputation and quality control, 1 146 individuals had genotype data, covariate data and phenotype data simultaneously (age: 115.25±9.59 months; sex ratio(M:F): 588:558) (Table S3 for sample size of each continuous trait analysed in this study).

We conducted GWASs with genotype dosage using PLINK v1.90. Linear regression analyses for 11 mathematical ability categories were carried out by an additive model using quadratic term for age, sex, school and the first five principal components of population structure as covariates [19]. Population structure analysis was conducted by PLINK v1.90 and GCTA. To perform this analysis, we used options of–autosome, make-grm, grm and pca in the GCTA. Quantile-quantile plots were generated by package qq-man (R3.5) to reveal the consistency between the observed and expected *P* value and the rationality of analysis model. Manhattan plots were generated by package qq-man (R3.5) to present the result of GWAS. Genome-wide significance was defined as common threshold (*p*=5×10^−8^). Since the 11 phenotypes are highly correlated, we did not correct for the number of phenotypes tested. Regional association plots were generated by a web-based tool LocusZoom (http://locuszoom.org/) to reveal regional visualizations around top SNPs. Analysis was performed on autosomes and unrelated individuals only. We also examined the impact of identified significant loci on other phenotypes.

### Assessment of SNPs previously associated with mathematical ability

To date, a total of 988 genome-wide significant (*p*<5×10^−8^) SNPs have been reported by previous publications. One of these SNPs (rs133885) associated with a composite score (addition, multiplication and numerosity judgements) was discovered in German-Austrian dyslexia samples (*p*=7.71×10^−10^, *n*=699) and replicated in a sample representing the general population (*p*=0.048, *n=1* 080) [11]. Four of these SNPs associated with Chinese students’ curriculum scores of mathematics were reported from a study of meta-analysis of results in three different cohorts, and these four SNPs have strong linkage disequilibrium [7]. The other 983 SNPs, of which 618 SNPs associated with self-reported math ability (*n*=564 698) and 365 SNPs associated with highest math class taken (*n*=430 445), were identified in samples from 23andMe, and all these SNPs are independent [10]. We looked up these SNPs in our data, mapped 585 SNPs, and checked whether they were associated with mathematical ability categories measured in Chinese children. Here we used a loose criterion *p* = 0.05/585 for significant replication. Details of these SNPs can be found in Table S4.

### Gene- and gene set-based enrichment tests

Gene- and gene-set-based enrichment analyses for the mathematical ability categories were performed by MAGMA [20]. The gene analysis was based on our raw GWAS data. First of all, SNPs were allocated on the protein-coding genes according to their genomic location. These genes containing at least one SNP of our data were then subjected to internal quality control. Finally, the filtered genes were tested in the gene-based enrichment analysis. Given the number of genes (17 734) finally tested, the Bonferroni-corrected significance level for this analysis was set to 2.82×10^−6^ (*p*=0.05/17 734).

The gene-set analysis used a competitive gene-set analysis in MAGMA based on the result of gene analysis [21]. Original gene sets are from C2 data set (including Chemical and genetic perturbations, Canonical pathways) on the Gene Set Enrichment Analysis. We tested a total of 5 497 gene sets containing genes defined in the genotype data. Given the number of gene sets finally analyzed, the Bonferroni-corrected significance level for this analysis was set to 9.10×10^−6^ (*p*=0.05/5 497).

## RESULTS

### Single-variant genome-wide associations

We separately analyzed eleven mathematical ability categories in this study. Mathematical reasoning ability showed genome-wide significant associations with seven SNPs, the most significant SNP we identified is rs34034296 (*p*=2.01×10^−8^, *MAF*=0.185). Because the seven SNPs were in high LD with each other (all r^2^>0.8), indeed we only had one independent hit associated with this trait (we report the most significant one). All these SNPs mapped to an intergenic region with the nearest gene CSMD3 about 400 kb away from them.

8

Spatial conception ability showed a nominally genome-wide significant association with rs1369404 (*p*=5.40×10^−8^, *MAF*=0.062).

Manhattan plots for mathematical reasoning and spatial conception abilities are revealed in Figure 1 and 2. Regional association plots with these two categories are revealed in Figure 3 and 4. More details for all significant SNPs are reported in Table 1. Quantile-quantile plots for all traits and Manhattan plots for other traits are reported in Supplementary Figures S1-20. Results of genome-wide association analysis of all phenotypes (p<10^−5^) are presented in supplementary Table S5.

**Figure 1.**
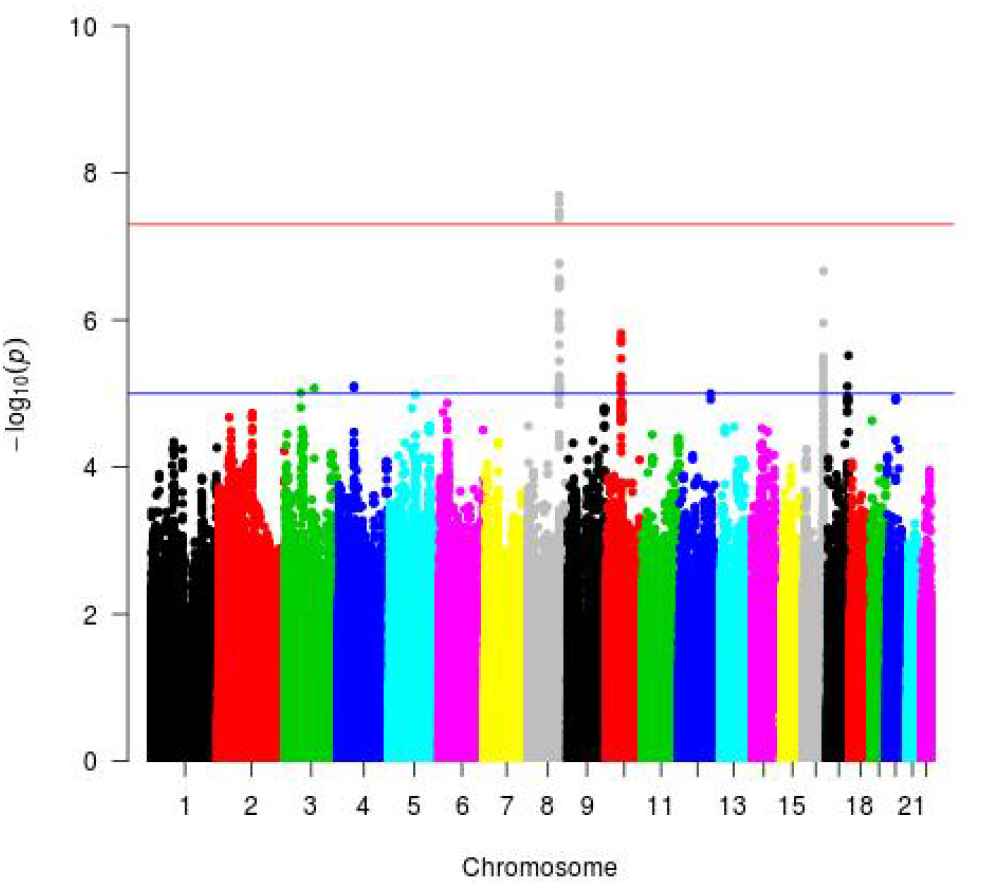
Manhattan plots for mathematical reasoning ability. X-axis represents the position of SNPs on each autosome, and y-axis represents the significance of association with the phenotype. The horizontal blue line represents a threshold of significance that we specified (1×10^−6^). The horizontal red line represents the threshold for genome-wide significance (5×10^−8^).

**Figure 2.**
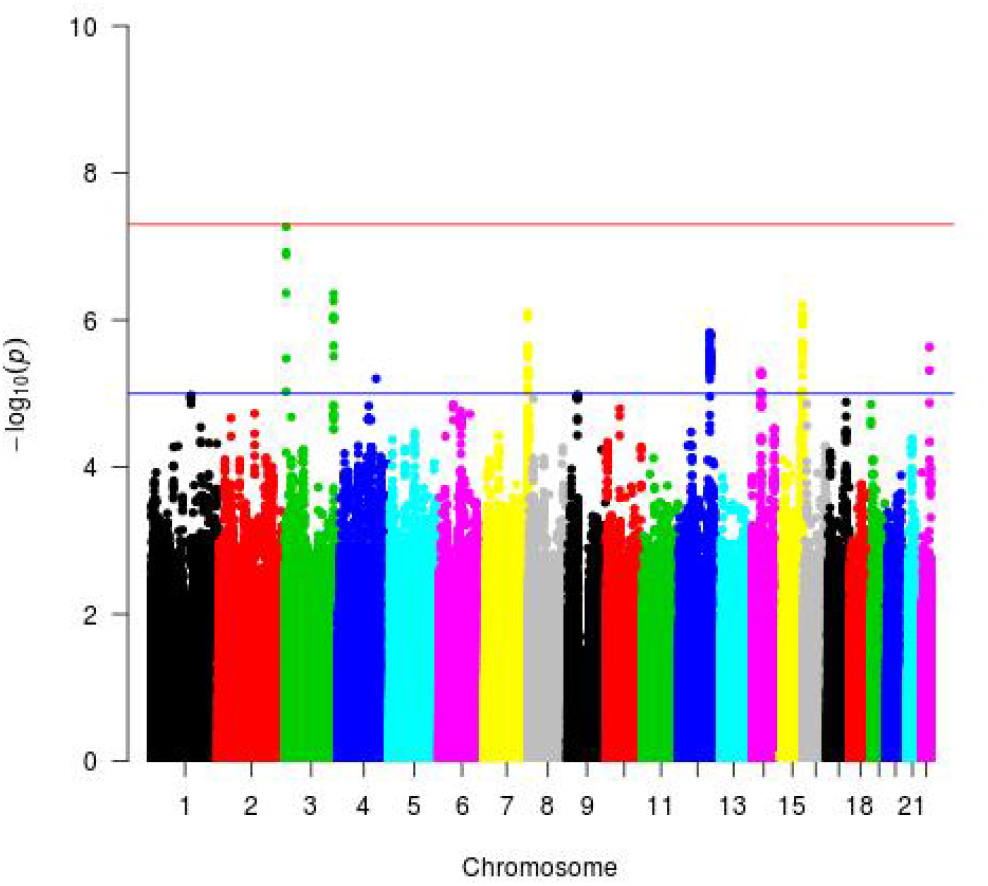
Manhattan plots for spatial conception ability. X-axis represents the position of SNPs on each autosome, and y-axis represents the significance of association with the phenotype. The horizontal blue line represents a threshold of significance that we specified (1×10^−6^). The horizontal red line represents the threshold for genome-wide significance (5×10^−8^).

**Figure 3.**
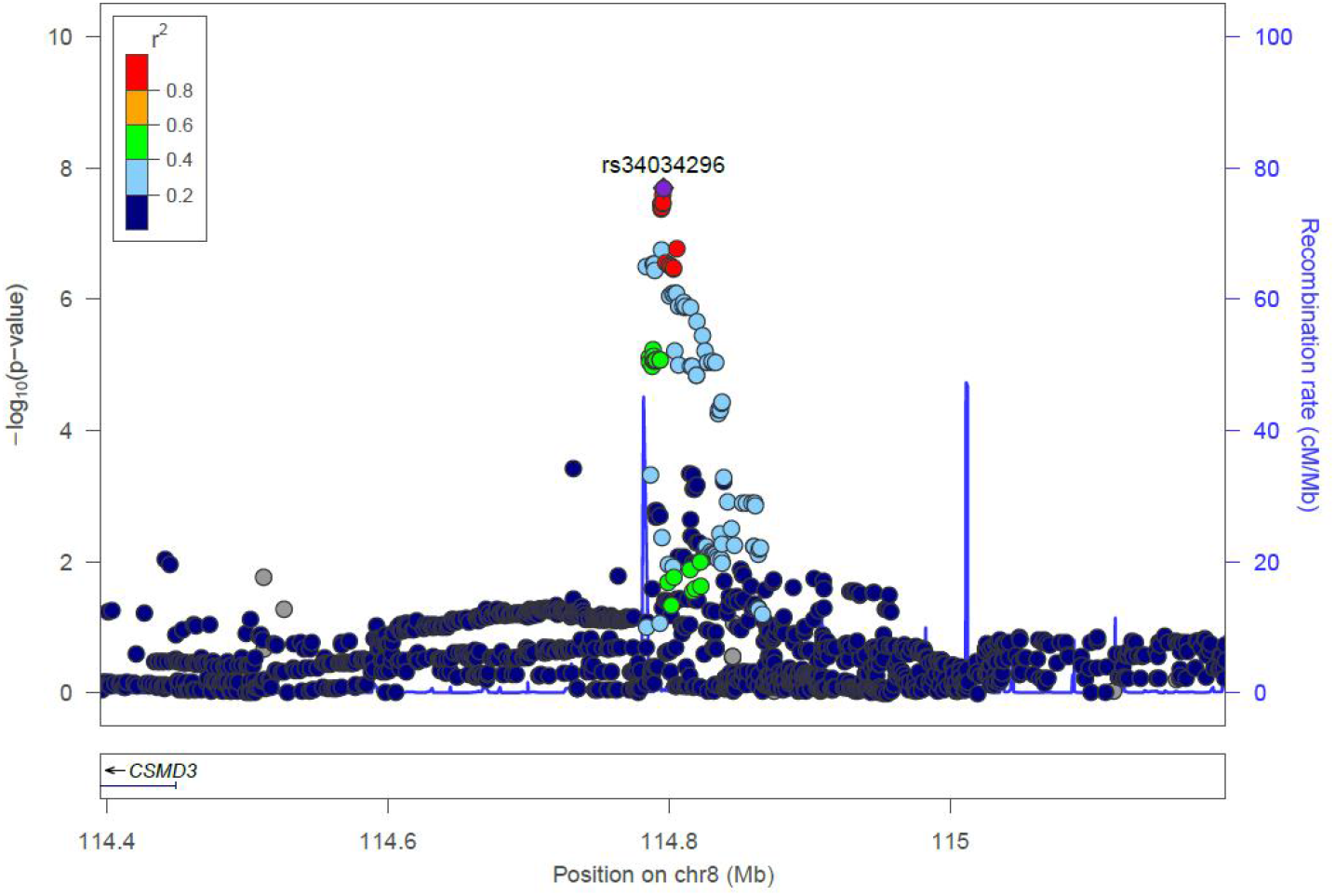
Regional association plots with mathematical reasoning ability. The most significant genetic loci are highlighted in violet. r^2^ are the LD values between other loci and the most significant SNP.

**Figure 4.**
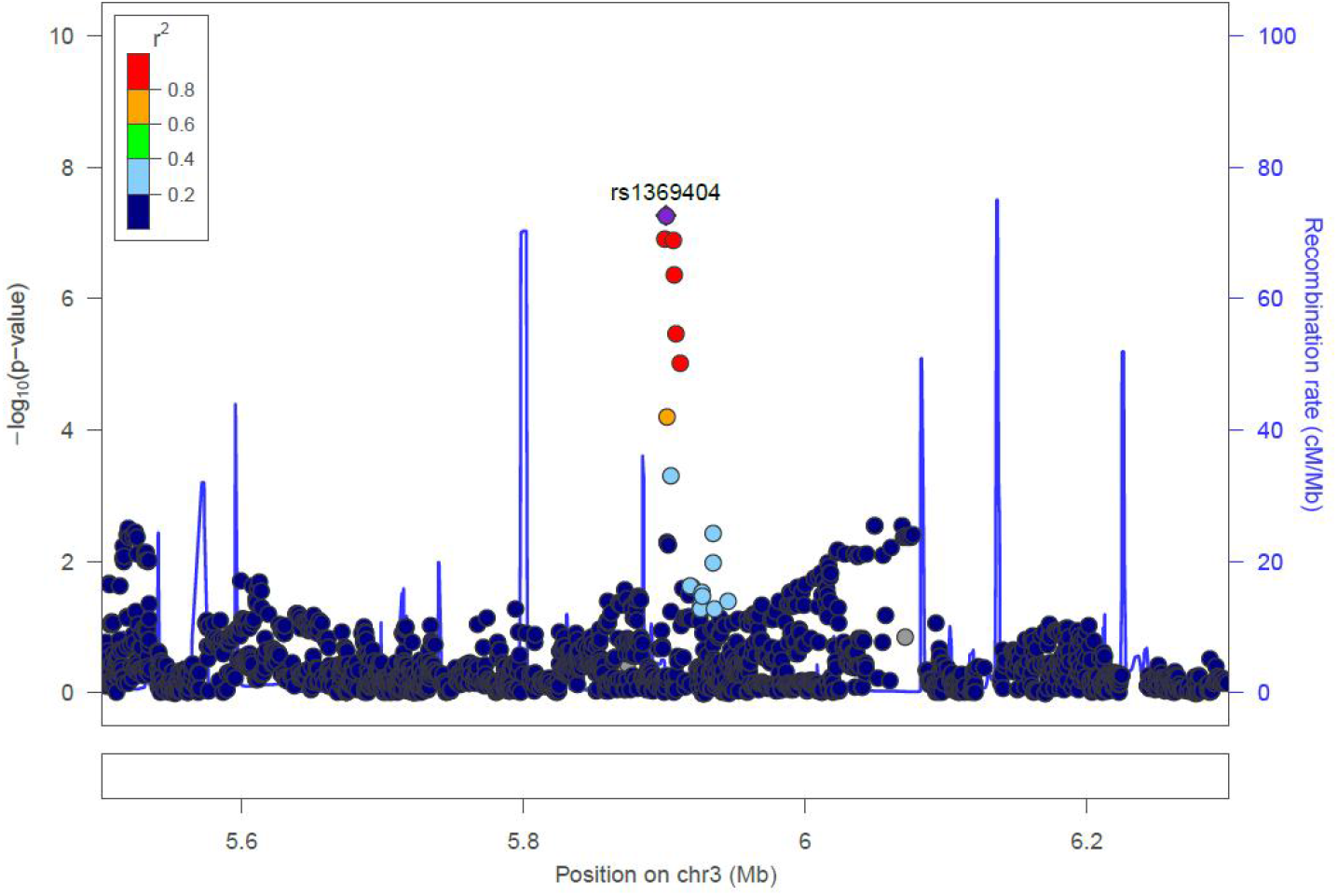
Regional association plots with spatial conception. The most significant genetic loci are highlighted in violet. r^2^ are the LD values between other loci and the most significant SNP.

**Table 1.**
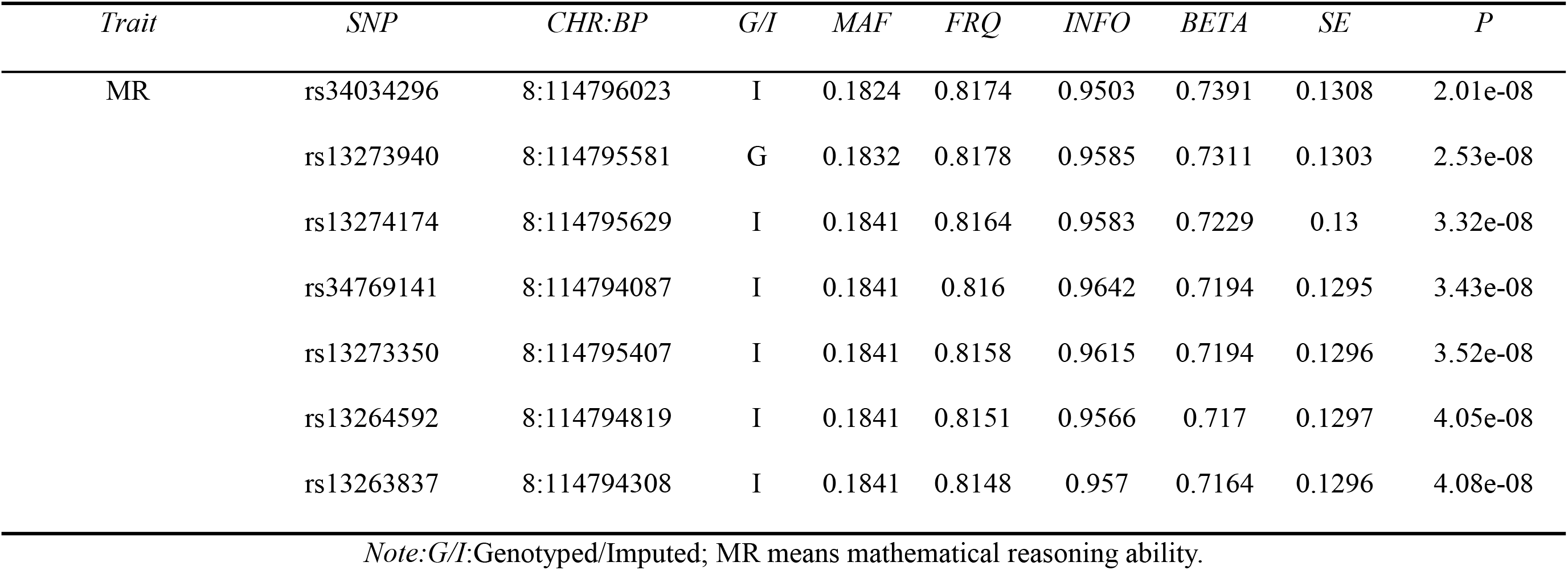
Seven significant genetic loci (*p*<5×10^−8^) detected in the GWAS analyses using genotype dosage.

The most significant SNP (rs34034296) also showed an association with another trait (magnitude perception) withstanding the Bonnferoni correction (*p*=2.638×10^−3^ < 0.05/10). Detailed information of analyses of the relationship between the SNP and all other traits are reported in Table 2.

**Table 2.**
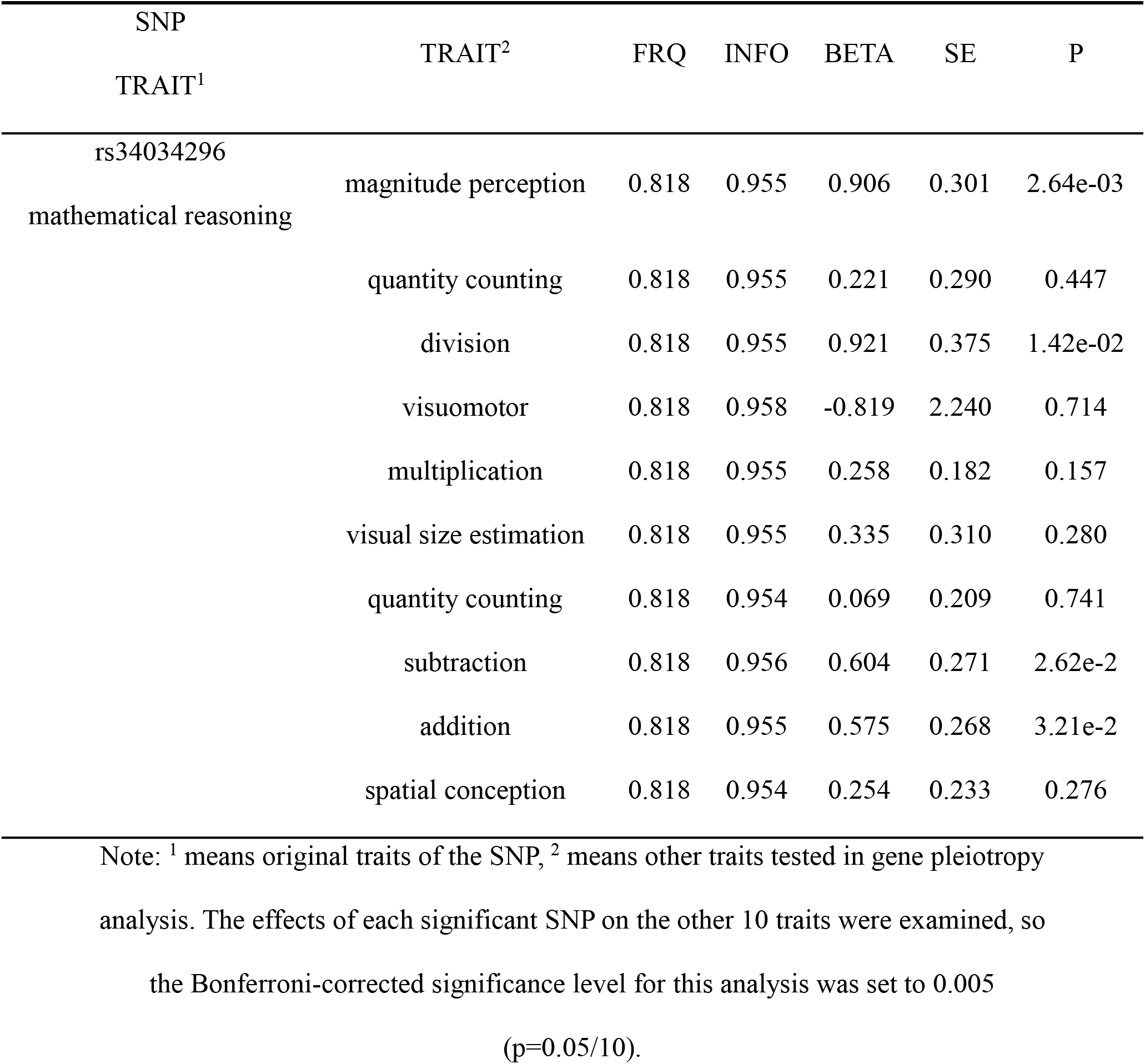
Results of pleiotropy for the significant SNP.

### SNPs previously reported to be associated with general mathematical ability

We replicated one genetic locus (rs133885) associated with division ability in our data (*p*=1.053×10^−5^) that survived Bonferroni-corrected significance level of 8.5×10^−5^ (*p*=0.05/585). The risk allele is identical in these two studies.

### Gene- and gene-set-based associations

We observed a significant association for a gene LINGO2 (leucine rich repeat and lg domain containing 2) with subtraction ability (*Z*=4.60,*p*=2.08×10^−6^), a significant association for a gene OAS1 (2’-5’-oligoadenylate synthetase 1) with spatial conception ability (*Z*=4.62, *p*=1.90×10^−6^), a significant association for a gene HECTD1 (HECT domain E3 ubiquitin protein ligase 1) with division ability (*Z*=4.54, *p*=2.80×10^−6^).

Similarly, in the gene-set-based analysis, we observed a significant association for a gene set (REACTOME_ERYTHROPOIETIN_ACTIVATES_PHOSPHOINOSITIDE_3_KINASE) with magnitude perception ability (*p*=3.50×10^−7^, *β*=1.36, *SE*=0.27), a significant association for a gene set (DASU_IL6_SIGNALING_DN) with addition ability (*p*=1.42×10^−6^, *β*=1.31, *SΛ*’ 0.28), a significant association for a gene set (DASU_IL6_SIGNALING_DN) with division ability (*p*=3.97×10^−8^, *β*=1.52, *SE*=0.28) and a significant association for a gene set (BIOCARTA_P53_PATHWAY) with spatial conception (*p*=1.88×10^−6^, *β*=0.80, *SE*=0.17). The genes in these three gene sets are listed in Table S6a-d.

## DISCUSSION

This is the first time that Heidelberg mathematics test has been used to measure phenotypes in GWAS of mathematical ability. We identified seven SNPs associated with mathematical reasoning ability. We replicated one SNP (rs133885) from the 585 SNPs previously reported to be associated with general mathematical ability. We also found genes or gene sets significantly associated with addition, subtraction, division, magnitude perception and spatial conception ability in enrichment analysis.

The most significant variant associated with mathematical reasoning ability was rs34034296. This SNP is located in the desert region of genome. The nearest gene to this locus is CSMD3. Researchers have reported copy number variants (CNVs) of CSMD3 in patients with schizophrenia and autism [22–24]. Autism and developmental dyscalculia are neurodevelopmental disorders, and they have comorbidities [25]. We for the first time show that these genes are directly associated with mathematical ability.

We assessed SNPs previously associated with mathematical ability. One SNP (rs133885) associated with a composite score (addition, multiplication and numerosity judgements) was replicated in the phenotype of division ability in our data. In multiplication task, children were asked to judge whether the presented equations (e.g. “5×6=30”) were correct. Division is the inverse of multiplication, so children may use division to test the correctness of multiplication equations. SNPs associated with self-reported math ability and highest math class taken were not replicated in any phenotype. Self-reported math ability and highest math class taken were assessed by questions, which is very different from the way we evaluated mathematical ability. Also, SNPs associated with a composite curriculum score (understanding numbers, computing and knowledge, non-numerical processes) were not replicated. Therefore, different ways and dimensions of evaluating mathematical ability may affect the replications of significant loci. Other reasons that significant loci were not replicated may be related to our relatively small sample size and ethnicity differences.

Among these identified genes and gene sets, there are two genes associated with cognition. LINGO2 was identified as a gene significantly associated with subtraction ability. It can regulate synapse assembly and was reported to be a risk gene for autism spectrum disorders [26]. OAS1 was identified as a gene significantly associated with spatial conception ability. It has ubiquitous expression in various tissues and was reported to be a risk gene for Alzheimer’s disease [27]. Similarly, these genes and gene sets above have been identified for the first time to be associated with mathematical ability.

In conclusion, we identified SNPs, genes and gene sets associated with mathematical skills in a sample of Chinese children. Results of our research provide evidence that different mathematical abilities may have different genetic basis. This study not only refined the current GWAS of mathematical ability but also added some population diversity to the literature. Future studies should expand the sample size to verify our findings.

## Acknowledgments

The study was funded by National Natural Science Foundation of China (61807023), Humanities and Social Science Fund of Ministry of Education of the People’s Republic of China (17XJC190010), Shaanxi Province Natural Science Foundation (2018JQ8015), and Fundamental Research Funds for the Central Universities (CN) (GK201702011) to Jingjing Zhao. This study was also supported by internal funds from Shaanxi Normal University to Jingjing Zhao and Chunyu Liu. The authors would like to thank all children who participated in the study and their parents and teachers for their time and cooperation.

## SUPPORTING INFORMATION

Additional supporting information may be found in the Supporting Information section at the end of this article at MP’s website.

