## Supplemental Materials for "A genome-wide association study identified new variants associated with mathematical abilities in Chinese children"

**Supplementary Information**

**Supplementary METHODS**

**Table S1. Mathematical ability categories of the test analyzed in the present study**

| **Categories** | **Examples of tasks** | **Introduction** |
| --- | --- | --- |
| addition | 3 + 1 = __ | addition calculation of varied difficulty |
| subtraction | 3 - 1 = __ | subtraction calculation of varied difficulty |
| multiplication | 3 * 3 = __ | multiplication calculation of varied difficulty |
| division | 3 ÷ 1 = __ | division calculation of varied difficulty |
| equation computation | __ + 5 = 10 – 1 | filling in the missing number of varied difficulty |
| magnitude perception | 6 (> <or =) 5 | writing “(> <or =)” to indicate which number is larger, or both sides are the same. |
| mathematical reasoning | 0 3 0 4 0 5 _ _ _ | filling in the missing number according to the rule of the previous numbers |
| visual size estimation | * | estimating the size of the line segment |
| quantity counting | * | counting the number of objects arranged in a certain way |
| spatial conception | * | determining the number of cubes in a construction |
| visuomotor | * | connecting the numbers presented in the correct order |

***Note:*** * The subtests represented by graph can't be represented by table. More information can be seen in Haffner’s book [1].

**Testing process of Heidelberg mathematics test:**

For each subtest, first of all, the teacher made an introduction by reading instruction word by word, then students had to complete several exercises correctly in order to fully understand the requirements of the task before the formal test, at last, students were asked to complete items as many as possible in the same limited time. All teachers received the same training by experimenters. We strictly followed the standard answer of the test to score.

**Table S2.** **Cross-trait** **correlations (Pearson’s *r* coefficient) of the eleven** **mathematical ability categories calculated in the present study.**

| **Trait** | addition | subtraction | MR | multiplication | division | equation | MP | VSE | QC | SC | VM |
| --- | --- | --- | --- | --- | --- | --- | --- | --- | --- | --- | --- |
| addition | 1 |  |  |  |  |  |  |  |  |  |  |
| subtraction | .730 | 1 |  |  |  |  |  |  |  |  |  |
| MR | .323 | .330 | 1 |  |  |  |  |  |  |  |  |
| multiplication | .615 | .544 | .254 | 1 |  |  |  |  |  |  |  |
| division | .699 | .660 | .339 | .641 | 1 |  |  |  |  |  |  |
| equation | .660 | .702 | .465 | .523 | .665 | 1 |  |  |  |  |  |
| MP | .567 | .542 | .273 | .484 | .521 | .538 | 1 |  |  |  |  |
| VSE | .322 | .307 | .340 | .270 | .336 | .439 | .292 | 1 |  |  |  |
| QC | .453 | .446 | .208 | .365 | .414 | .384 | .548 | .264 | 1 |  |  |
| SC | .313 | .316 | .283 | .190 | .237 | .401 | .309 | .384 | .321 | 1 |  |
| VM | .260 | .196 | .133 | .205 | .226 | .200 | .329 | .106 | .319 | .133 | 1 |

***Note:*** MR means mathematical reasoning ability, MP means magnitude perception ability, VSE means visual size estimation ability, QC means quantity counting ability, SC means spatial conception ability, VM means visuomotor skill.

**Table S3. sample size of each continuous trait analysed in this study**

| **Trait** | addition | subtraction | MR | multiplication | division | equation | MP | VSE | QC | SC | VM |
| --- | --- | --- | --- | --- | --- | --- | --- | --- | --- | --- | --- |
| N | 1143 | 1144 | 1134 | 1143 | 1145 | 1145 | 1145 | 1143 | 1144 | 1144 | 1127 |

***Note:*** The traits of abbreviations in the above table are the same as those in Table S2.

**Supplementary RESULTS**

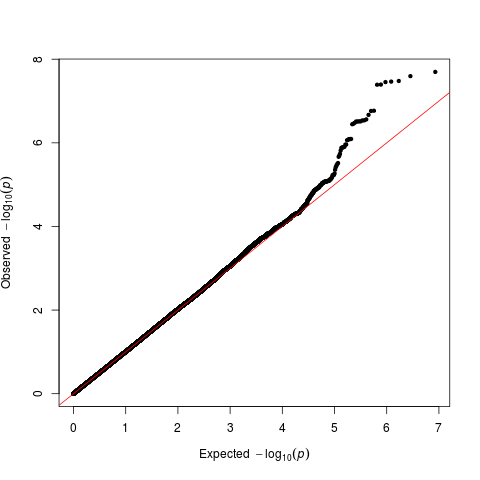

**Figure S1. Q–Q plot for mathematical reasoning ability (λ=** **1.006).**

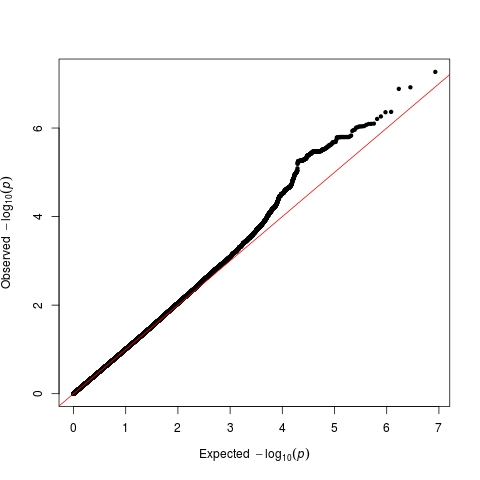

**Figure S2. Q–Q plot for spatial conception ability (λ=** **1.003).**

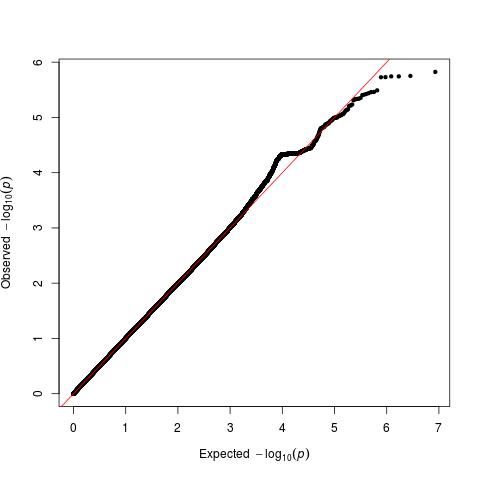

**Figure S3. Q–Q plot for** **addition power (λ=** **1.011).**

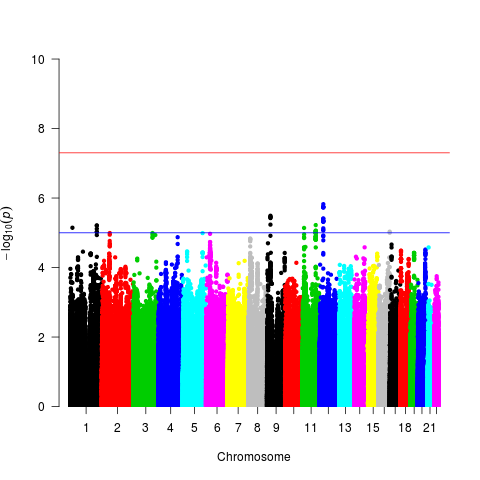

**Figure S4. Manhattan plot for addition power.**

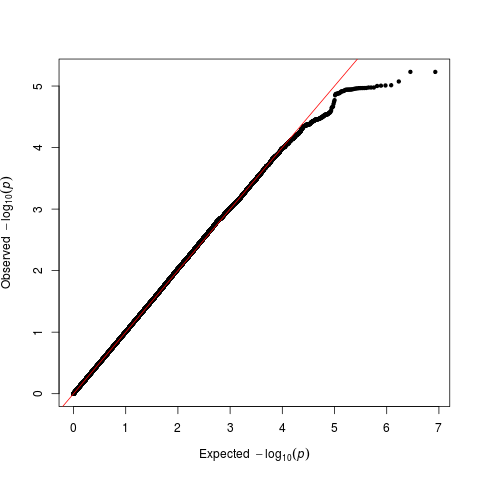

**Figure S5. Q–Q plot for subtraction power (λ=** **1).**

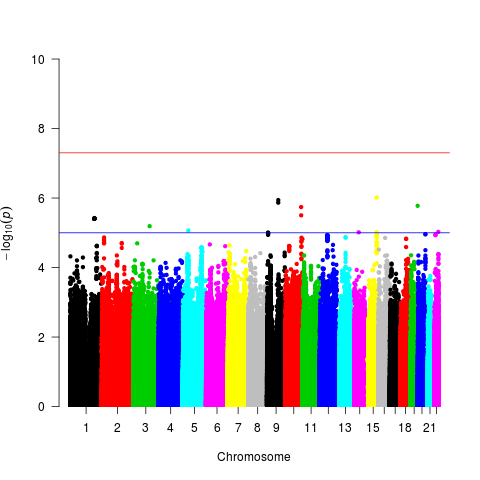

**Figure S6.** **Manhattan plot for subtraction power.**

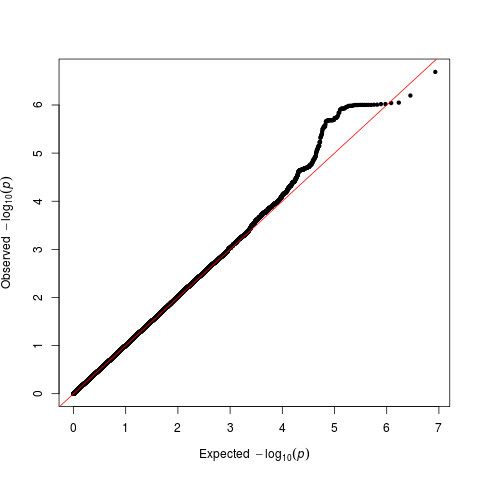

**Figure S7. Q–Q plot for multiplication power (λ=** **1.003).**

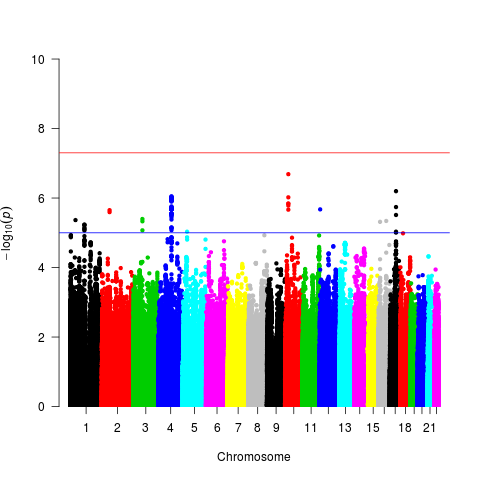

**Figure S8. Manhattan plot for** **multiplication power.**

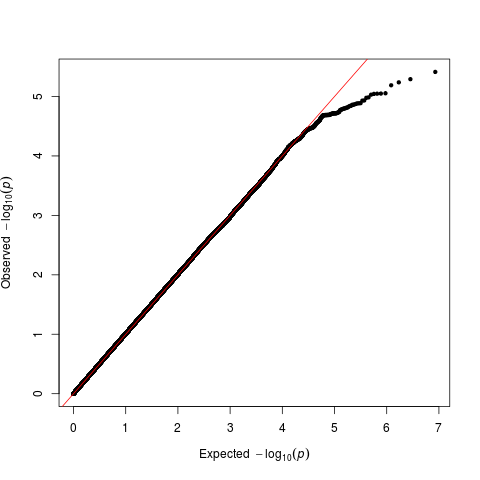

**Figure S9. Q–Q plot for division power (λ=** **1.018).**

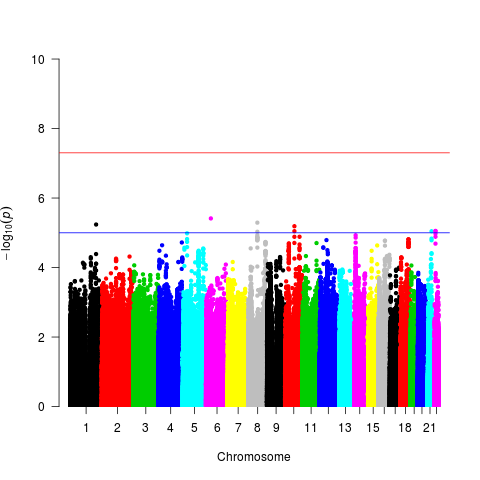

**Figure S10. Manhattan plot for division power.**

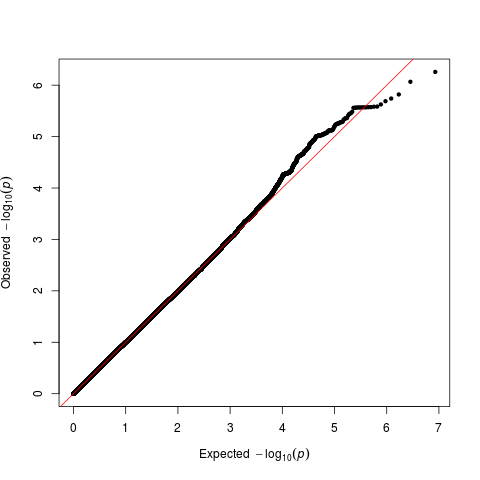

**Figure S11. Q–Q plot for equation power (λ=** **1.003).**

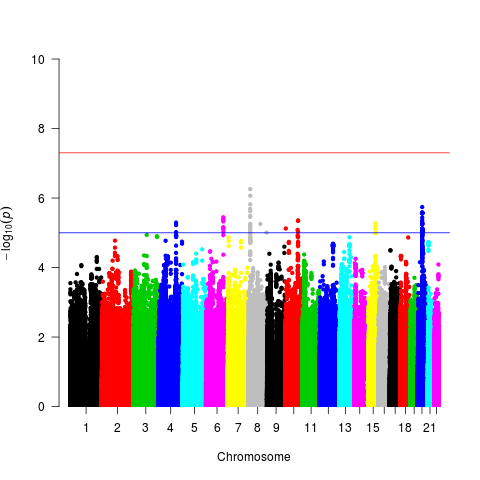

**Figure S12. Manhattan plot for equation power.**

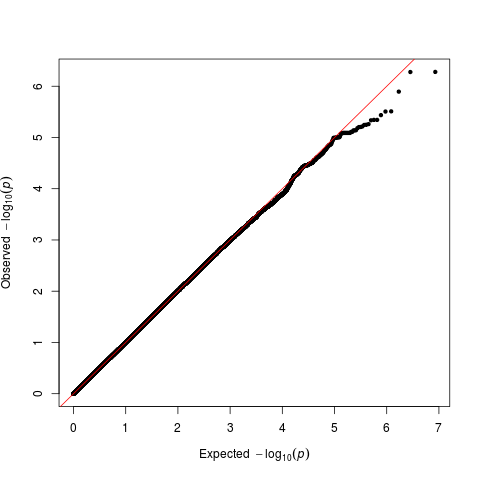

**Figure S13. Q–Q plot for** **magnitude perception ability (λ=** **0.996).**

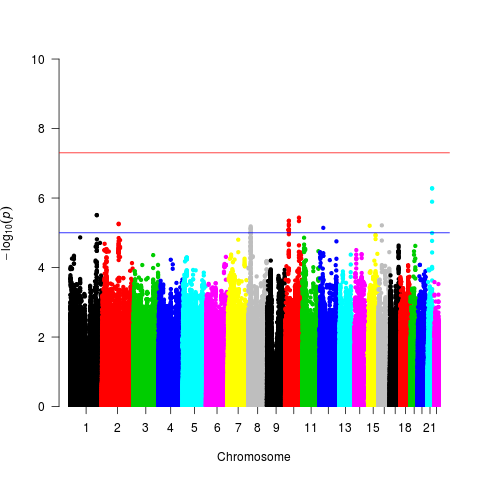

**Figure S14. Manhattan plot for magnitude perception ability.**

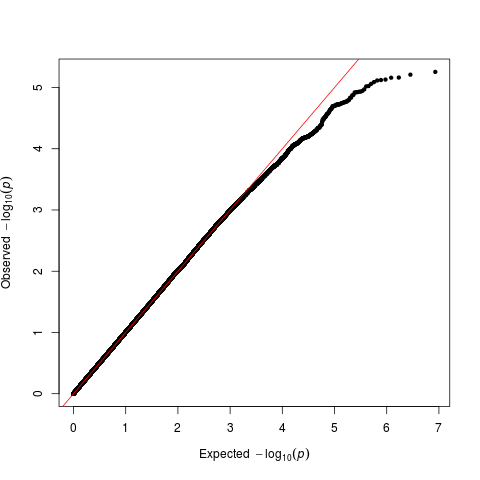

**Figure S15. Q–Q plot for** **visual size estimation ability (λ=** **1.016).**

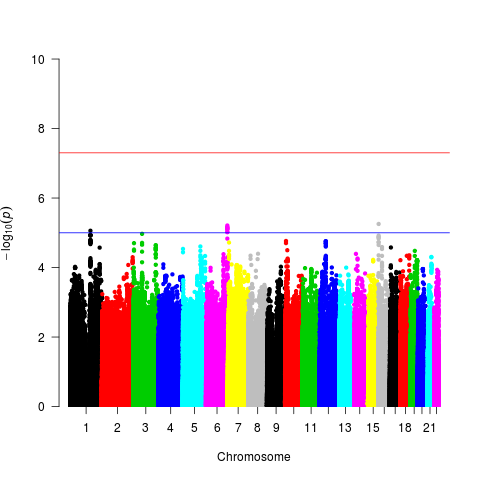

**Figure S16. Manhattan plot for visual size estimation ability.**

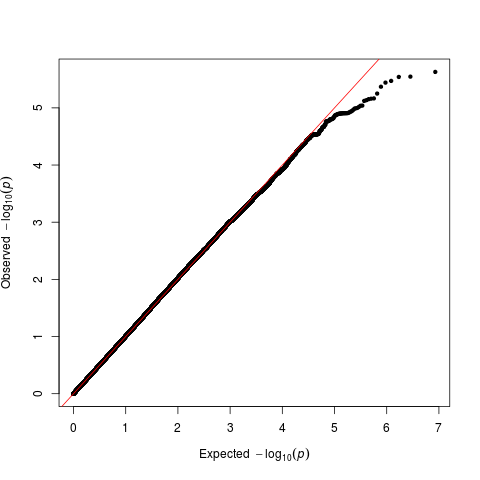

**Figure S17. Q–Q plot for** **quantity counting ability (λ=** **1.001).**

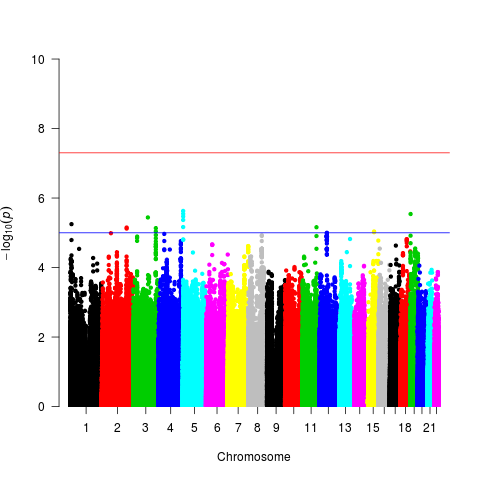

**Figure S18. Manhattan plot for quantity counting ability.**

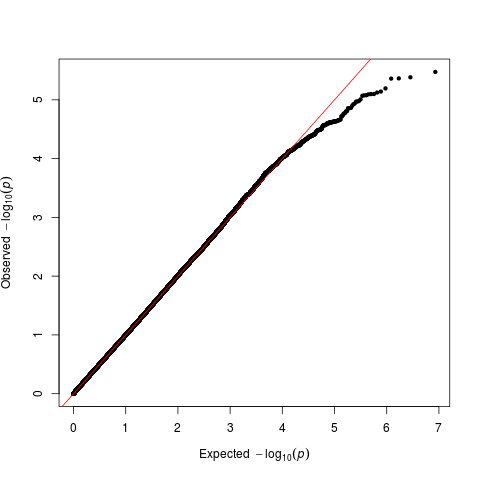

**Figure S19. Q–Q plot for visuomotor skill (λ=** **1.014).**

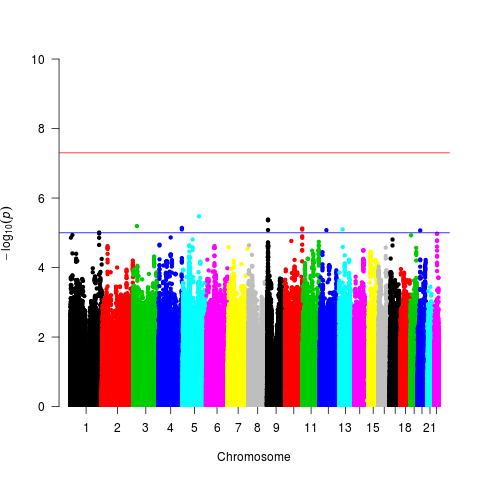

**Figure S20. Manhattan plot for visuomotor skill**

**Table S6a. gene list of set1 (REACTOME_ERYTHROPOIETIN_ACTIVATES_PHOSPHOINOSITIDE_3_KINASE_PI3K) (*P*= 3.50145e-07)**

| **GENE** | **CHR** | **START** | **STOP** | **NSNPS** | **NPARAM** | **N** | **ZSTAT** | **P** | **ZFITTED_BASE** | **ZRESID_BASE** |
| --- | --- | --- | --- | --- | --- | --- | --- | --- | --- | --- |
| 5293 | 1 | 9711789 | 9789172 | 124 | 15 | 1000 | 0.098004 | 0.47002 | -2.92E-05 | 0.098034 |
| 5291 | 3 | 138371540 | 138553780 | 26 | 7 | 1000 | 1.3478 | 0.090553 | -2.92E-05 | 1.3478 |
| 5290 | 3 | 178866311 | 178952500 | 100 | 6 | 1000 | 0.88091 | 0.19035 | -2.92E-05 | 0.88094 |
| 2549 | 4 | 144257680 | 144395718 | 95 | 5 | 1000 | 1.6876 | 0.044719 | -2.92E-05 | 1.6876 |
| 5295 | 5 | 67511584 | 67597649 | 120 | 16 | 1000 | 2.0471 | 0.021439 | -2.92E-05 | 2.0472 |
| 2056 | 7 | 100318423 | 100321323 | 2 | 1 | 1000 | 2.0302 | 0.020534 | -2.92E-05 | 2.0302 |
| 5294 | 7 | 106505723 | 106547592 | 30 | 4 | 1000 | 0.053799 | 0.4743 | -2.92E-05 | 0.053828 |
| 4067 | 8 | 56792386 | 56925006 | 82 | 8 | 1000 | 1.6943 | 0.045609 | -2.92E-05 | 1.6943 |
| 3717 | 9 | 4985245 | 5129948 | 242 | 9 | 1000 | 1.6039 | 0.05308 | -2.92E-05 | 1.604 |
| 8660 | 13 | 110406184 | 110438914 | 57 | 7 | 1000 | 1.5627 | 0.059226 | -2.92E-05 | 1.5627 |
| 23533 | 17 | 8782233 | 8869029 | 90 | 10 | 1000 | 0.19378 | 0.42578 | -2.92E-05 | 0.19381 |

***Note:*** GENE, the ID corresponding to the gene in the referenced annotation file; CHR, the chromosome on which the gene is located; START/STOP, the boundaries on both sides of the gene; NSNPS, the number of SNPs mapped to the gene that were found in raw GWAS data and were not removed based on internal SNP QC; NPARAM, the number of parameters used in the model; N, the assumed sample size; ZSTAT, the Z-value of the gene; P, the gene p-value; ZFITTED_BASE and ZRESID_BASE, the fitted values and residual values for the base model to be used for post-hoc outlier checks.

**Table S6b. gene list of set2 (DASU_IL6_SIGNALING_DN) (P= 1.42027e-06)**

| **GENE** | **CHR** | **START** | **STOP** | **NSNPS** | **NPARAM** | **N** | **ZSTAT** | **P** | **ZFITTED_BASE** | **ZRESID_BASE** |
| --- | --- | --- | --- | --- | --- | --- | --- | --- | --- | --- |
| 55752 | 4 | 77870895 | 77961307 | 177 | 11 | 1000 | 0.94483 | 0.16783 | -2.33E-05 | 0.94485 |
| 440 | 7 | 97481429 | 97501854 | 46 | 8 | 1000 | 0.49383 | 0.302 | -2.33E-05 | 0.49386 |
| 5054 | 7 | 100770370 | 1.01E+08 | 19 | 2 | 1000 | 0.46366 | 0.31245 | -2.33E-05 | 0.46369 |
| 23213 | 8 | 70378859 | 70573147 | 116 | 26 | 1000 | 2.3559 | 0.009111 | -2.33E-05 | 2.3559 |
| 5507 | 10 | 93388197 | 93392858 | 1 | 1 | 1000 | -0.1843 | 0.5781 | -2.33E-05 | -0.18427 |
| 7009 | 12 | 50135293 | 50158717 | 18 | 2 | 1000 | 0.3763 | 0.35092 | -2.33E-05 | 0.37632 |
| 7168 | 15 | 63334838 | 63364114 | 30 | 5 | 1000 | 1.515 | 0.061448 | -2.33E-05 | 1.5151 |

**Table S6c. gene list of set3 (DASU_IL6_SIGNALING_DN) (*P*= 3.97328e-08)**

| **GENE** | **CHR** | **START** | **STOP** | **NSNPS** | **NPARAM** | **N** | **ZSTAT** | **P** | **ZFITTED_BASE** | **ZRESID_BASE** |
| --- | --- | --- | --- | --- | --- | --- | --- | --- | --- | --- |
| 55752 | 4 | 77870895 | 77961307 | 177 | 11 | 1000 | 0.61069 | 0.26646 | -1.48E-05 | 0.61071 |
| 440 | 7 | 97481429 | 97501854 | 46 | 8 | 1000 | 1.6056 | 0.054628 | -1.48E-05 | 1.6056 |
| 5054 | 7 | 100770370 | 100782547 | 19 | 2 | 1000 | 0.14919 | 0.43522 | -1.48E-05 | 0.1492 |
| 23213 | 8 | 70378859 | 70573147 | 116 | 26 | 1000 | 3.0845 | 0.001149 | -1.48E-05 | 3.0846 |
| 5507 | 10 | 93388197 | 93392858 | 1 | 1 | 1000 | -0.80642 | 0.7881 | -1.48E-05 | -0.8064 |
| 7009 | 12 | 50135293 | 50158717 | 18 | 2 | 1000 | 0.42572 | 0.32354 | -1.48E-05 | 0.42573 |
| 7168 | 15 | 63334838 | 63364114 | 30 | 5 | 1000 | 1.0032 | 0.15786 | -1.48E-05 | 1.0032 |

**Table S6d. gene list of set3 (BIOCARTA_P53_PATHWAY) (*P*= 1.88201e-06)**

| **GENE** | **CHR** | **START** | **STOP** | **NSNPS** | **NPARAM** | **N** | **ZSTAT** | **P** | **ZFITTED_BASE** | **ZRESID_BASE** |
| --- | --- | --- | --- | --- | --- | --- | --- | --- | --- | --- |
| 1647 | 1 | 68150860 | 68154021 | 5 | 2 | 1000 | -0.24409 | 0.59842 | -6.81E-06 | -0.24409 |
| 1026 | 6 | 36644237 | 36655116 | 28 | 6 | 1000 | -0.9346 | 0.83028 | -6.81E-06 | -0.93459 |
| 595 | 11 | 69455873 | 69469242 | 9 | 2 | 1000 | 1.0887 | 0.13621 | -6.81E-06 | 1.0887 |
| 472 | 11 | 108093559 | 108239829 | 108 | 2 | 1000 | 1.206 | 0.10964 | -6.81E-06 | 1.206 |
| 1017 | 12 | 56360556 | 56366573 | 3 | 1 | 1000 | 2.9694 | 0.0014 | -6.81E-06 | 2.9694 |
| 1019 | 12 | 58141510 | 58146304 | 4 | 2 | 1000 | -0.082779 | 0.53413 | -6.81E-06 | -0.082772 |
| 4193 | 12 | 69201952 | 69239324 | 40 | 3 | 1000 | 0.93654 | 0.17408 | -6.81E-06 | 0.93655 |
| 317 | 12 | 99039078 | 99129211 | 120 | 9 | 1000 | 0.38152 | 0.35612 | -6.81E-06 | 0.38153 |
| 5925 | 13 | 48877883 | 49056026 | 134 | 7 | 1000 | 0.36345 | 0.35754 | -6.81E-06 | 0.36345 |
| 7157 | 17 | 7571720 | 7590868 | 30 | 5 | 1000 | -0.55286 | 0.71421 | -6.81E-06 | -0.55285 |
| 596 | 18 | 60790579 | 60987011 | 284 | 42 | 1000 | 0.36718 | 0.37885 | -6.81E-06 | 0.36719 |
| 898 | 19 | 30302805 | 30315224 | 22 | 4 | 1000 | -0.77435 | 0.78277 | -6.81E-06 | -0.77434 |
| 581 | 19 | 49458117 | 49465055 | 1 | 1 | 1000 | 0.4093 | 0.3333 | -6.81E-06 | 0.4093 |
| 5111 | 20 | 5095599 | 5107268 | 1 | 1 | 1000 | 1.6012 | 0.04935 | -6.81E-06 | 1.6012 |
| 1869 | 20 | 32263292 | 32274210 | 5 | 1 | 1000 | -0.56025 | 0.70828 | -6.81E-06 | -0.56024 |
| 7078 | 22 | 33196802 | 33259028 | 74 | 11 | 1000 | 1.2709 | 0.10714 | -6.81E-06 | 1.2709 |
